# A Self-Purifying Microfluidic System for Identifying Drugs Acting Against Adult Schistosomes

**DOI:** 10.1101/2022.04.04.486714

**Authors:** Vincent Girod, Marie-José Ghoris, Stéphanie Caby, Oleg Melnyk, Colette Dissous, Vincent Senez, Jérôme Vicogne

## Abstract

The discovery of novel antihelminthic molecules to combat the development and spread of schistosomiasis, a disease caused by several Schistosoma flatworm species, mobilizes significant research efforts worldwide. In the absence of reliable and practical biochemical assays for measuring the viability of adult worms, the antischistosomicidal activity of molecules is usually evaluated by a detailed microscopic observation of worm mobility and/or integrity upon drug exposure. These assays have the disadvantage of being inacurate, subjective, biased by the limited in vitro worm viability and difficult to integrate at high density. We describe here a self-purifiying microfluidic system enabling the selection of healthy adult worms and the identification of molecules acting on the parasite. The worms are assayed in a dynamic environment that eliminates unhealthy worms that cannot attach firmly to the chip walls prior to being exposed to the drug. The detachment of the worms is also used as second step readout for identifying active compounds. We have validated this new fluidic screening approach using the two major antihelmintic drugs, Praziquantel and Artemisinin. The reported dynamic system is simple to produce and to parallelize. Importantly, it enables a quick, sensitive and reliable detection of antischistosomal compounds in no more than one day. This system can potentially be modified in the future to better mimic the natural habitat of the parasite.

## Introduction

Schistosomiasis is caused by flatworms of the genus Schistosoma and is the second most important parasitic disease, infecting more than 240 million people worldwide^1–2^. An outbreak of schistosomiasis in Corsica (France) since 2013 raises fears of an expansion of this parasitosis in southern Europe in the coming decades^3^. The pathology of schistosomiasis is due to the reaction of the host to schistosome eggs that accumulate in tissues during their intense and permanent production by adult worm couples, a problem worsened by their remarkable longevity (several decades). In the mammal host, mature male and female worms are paired and reside in blood vessels in a final location that varies according to the schistosome species. Adult couples of *Schistosoma mansoni*, the species that shows the widest geographical distribution and used in this study, locate to the mesenteric veins (Fig. 1a)^4^.

**Fig. 1:**
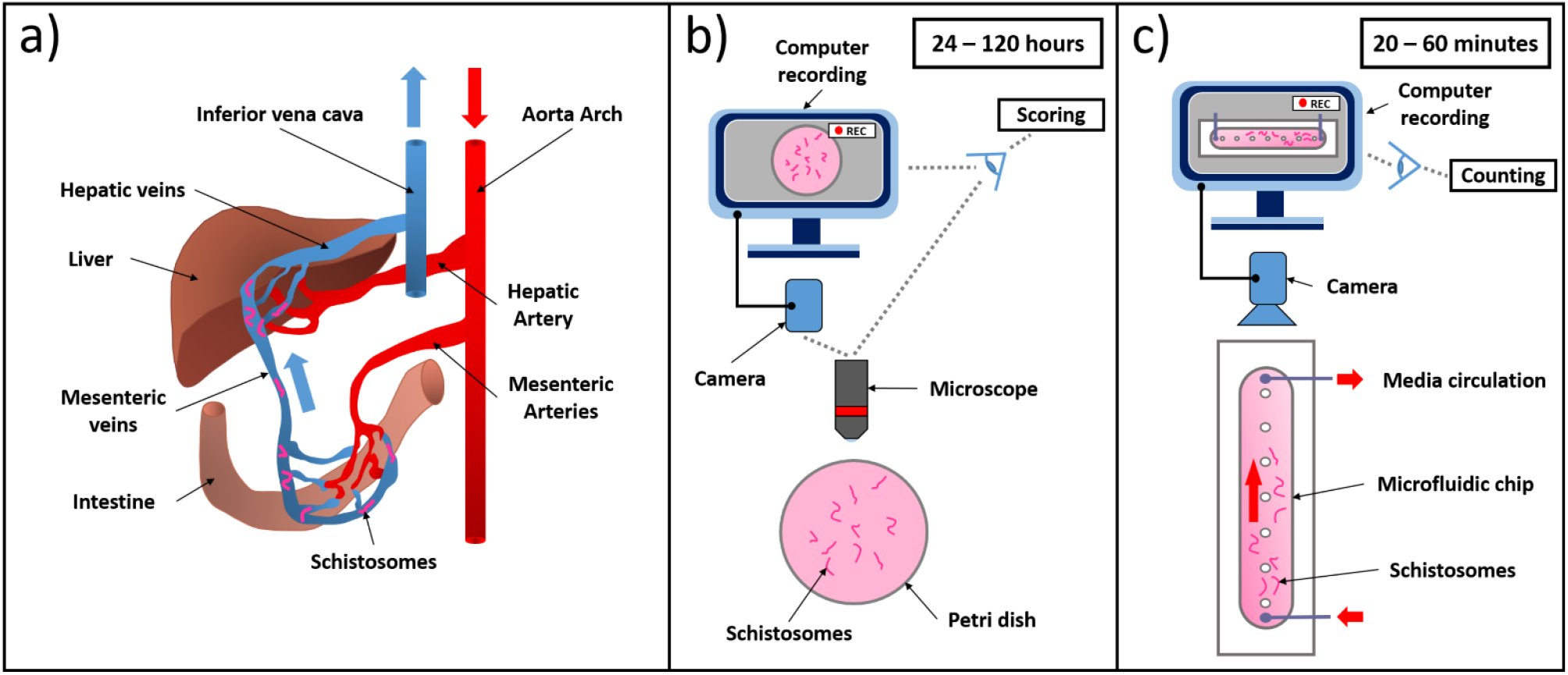
Physiological localisation of schistosomes in vertebrate hosts and comparation of static and dynamic drug characterisation studies. a) In their mammal hosts, adult schistosomes are localized in the mesenteric veins with bloodstream oriented from intestine towards the liver. b) During in vitro studies, worms are cultivated in static conditions in Petri dishes and observed under the microscope for 24 to 120 hours to analyse phenotypes and survival rates. c) In the dynamic system reported here, worms are introduced in a microfluidic chip with an oriented flow that force them to grip to the wall surface. Worm number is determined for 20 to 60 min after the introduction of the drug.

Since there is a real difficulty in developing a vaccine against schistosomiasis^5^, the therapeutic treatment of the disease is still achieved by oral administration of praziquantel (PZQ)^6^. PZQ has a strong lethal effect against adult worms. Unfortunately, it is significantly less active against the juvenile forms that penetrated the skin, i.e. the schistosomula. Therefore, the treatment must be repeated regularly^7^. In addition, as PZQ has been used extensively in infected countries for many years, the development of tolerant parasite strains is now observed^8^. Specifically, lower efficacy has been demonstrated both in the laboratory and the field^9–11^. Therefore, there is now an urgent need to introduce new anti-schistosome treatments.

*S. mansoni* schistosomula, the juvenile form of the parasite can be produced in high numbers using fresh cercariae released from its intermediate host, the snail *Biomphalaria glabrata*. However, in absence of any *in vitro* culture system enabling mature worm production, adults have to be collected by blood perfusion from the mammal host, usually a rodent (mouse or hamster) and are assayed subsequently.

Different approaches to evaluate compounds against juvenile or mature worms have been developed^12^. Innovative assays have been devised for the analysis of schistosomula using image based^13–14^, metabolic^15–16^ and electro-impedance techniques^17^. Importantly, several of them can be automated and are able to assay a large number of juvenile worms, generating more relevant data, although they give no information about the efficacy of the compounds on the adult stage.

Comparatively, the assays analyzing the response of adult worms to drugs are less diversified and enable testing of only a limited number of worms (1-5 couples). None of these methods yet constitutes a gold standard assay^18^ despite the development of sophisticated image-^19^ and non-image-based analysis combining electrical impedance^20^ or heat flow^21^ parameters. Phenotypic analysis of whole worms *in vitro* still remains the most widely used approach to identify active compounds^22–23^. This is done by scoring morphological and/or behavioral changes, mostly centered on motility parameters (Fig. 1b)^24^. A numerical scale called the ‘severity score’^25^, usually comprising four^15^ or five^26^ score levels, is used to describe deleterious phenotypes. Although these scoring methods have been shown to be effective in identifying drug candidates that later showed efficacy in a mouse model of *S. mansoni* infection^27^, they are difficult to set up and parallelize due to the need for a manual assessment. Moreover, the limited viability of the worms *ex vivo* affects the reproducibility and reliability of these assays. Above all, these *in vitro* assays do not allow measurement of the effect of a drug whilst also taking into account the diminution of the viability of the worms over time, a problem that is worsened by the duration of phenotypic assays that are run for several days.

We therefore searched for a novel assay that would i) furnish a quantitative response to a drug within a few hours and not days, ii) be self-purifying by enabling the elimination of dying or weak worms before applying the drug.

We reasoned that assaying the worms under flow conditions might provide a solution to the above limitations. Indeed, within infected hosts, healthy worms continuously grip the wall of the veins to resist the blood flow. Depending on vessel flow, viscosity and geometrical constraints, adult worms either use their oral and ventral suckers or favor a crawling mode of locomotion^28^. A worm with compromised viability quickly loses its ability to grip and maintain its position against the flow. Therefore, the adhesion of worms under flow conditions might not only offer a simple and efficient means for selecting the worms before starting the biological assay, it might also provide a quantitative readout for measuring the effect of a drug (Fig. 1c).

In this work, we designed a single channel microfluidic chip able to host 30 couples. We have evaluated different surface coatings for their effect on promoting worm adhesion and motility under flow conditions. Having achieved the conditions for enabling the worms to strongly grip the chip walls by oral and ventral suckers, we used the chip to select healthy worms and probe their resistance to increasing doses of PZQ or Artemisinin (ART).

## Materials and methods

### Parasite material

All animal experimentations were conducted in accordance with the European Convention for the Protection of Vertebrate Animals used for Experimental and other Scientific Purposes (ETS No 123, revised Appendix A) and were approved by the local committee for ethics in animal experimentation (authorization No. APAFIS#8289-2016122015127050v3) and the Pasteur Institute of Lille (Agreement No. B59350009).

A Puerto Rican strain of *S. mansoni* is maintained in the laboratory using the intermediate snail host *Biomphalaria glabrata* and the definitive golden hamster host *Mesocricetus auratus. S. mansoni* adult worms were obtained by hepatic portal perfusion of hamsters, six weeks after having been infected by cercariae released from infected snails^29^. Perfused worms were immediately washed with RPMI culture medium (RPMI 1640 GIBCO, 61870-010, UK), transferred into fresh RPMI culture medium supplemented with heat-inactivated horse serum (10%, GIBCO, 16050-122, UK), rifampicin (60 μg.mL^−1^, EUROMEDEX, 1059, Souffelweyersheim, France), penicillin (50 units.mL^−1^) and streptomycin (50 μg.mL^−1^, SIGMA, P-4333, USA). This medium is called RPMI Horse hereinafter.

### Worm culture medium optimization

Typically, immediately after perfusion, 100-150 worms were incubated in 100 mm Petri dish plates containing 25 mL of complete RPMI medium for 2-3 h in order to allow separated male and female worms to pair again (37 °C, saturated humid atmosphere, 5% CO_2_). Worms were maintained for a maximum of 5 days after perfusion and culture medium was renewed every 12 h. For medium testing experiments, pools of 10 couples (n=8) were maintained up to 1 month in 6 well plates (9.6 cm^2^) using 3 mL of M199 (GIBCO, 41150-020, UK), DMEM (GIBCO, 61965-026, UK) or RPMI supplemented either with 10% of heat-inactivated fetal calf serum (PAA, A15-101, AT), horse serum or human serum (Institute Pasteur of Lille, FR). All medium/serum combinations were supplemented with antibiotics, as described above and 2 mL of medium were exchanged with fresh medium every 48 h.

### MDCK cell culture

Madin-Darby Canine Kidney cells (MDCK, ATCC, USA, p11, 2008) were cultured in DMEM medium supplemented with 10% of heat-inactivated fetal bovine serum (Gibco, 10270-106, USA), 1% (v/v) of non-essential amino acids (Gibco, 11140-050, USA), biotin (50 μg.mL^−1^, Sigma, B4639, USA) and ZellSchield^®^ 1X (ZellSchield®, 13-0150, Germany), called MDCK medium. At subconfluence, cells were seeded as advised according to the ATCC protocol.

### Analysis of worm viability

The viability of the worms was evaluated by observing the parasites with a macroscope (OPTIKA, SZ6745TR, Italy) and using a well-established motion status protocol^18,30^. We have defined a simplified ‘severity score’ based on 3 levels: S2 score is for worms having a normal movement and normal tissues and tegument aspects, S1 score corresponds to worms with reduced movement or degraded tissues or tegument, while S0 corresponds to dead worms having no movement or with strong tissue degradation. RStudio (1.1.463 release, RStudio Inc., Boston, USA) was used to correct data when an S0 score (dead) was followed by an S1 or S2 score (still alive) and thus to avoid falsely counting worms that were still alive as dead. The data are presented as percentages, mean values and standard deviations.

### Microfluidic chip design

The geometrical features of the chips were designed by using COMSOL MULTIPHYSICS (Comsol AB, Stockholm, Sweden) to obtain a single microfluidic chamber The fluid flow was calculated with the microfluidic module using laminar flow conditions assuming a very low Reynolds number (see supplementary Table S3). The 3D FEM model is made of about 2 106 192 degrees of freedom using the predefined ‘extra fine’ mesh refinement. In the fluid flow model, non-slip initial condition was imposed to all surfaces corresponding to a solid/liquid interface. A fixed flow rate (between 0.05 and 3.4 ml.min^−1^) was used for the inlet and a fixed pressure (P = 0 Pa) for the outlet. The equations were solved in 900 s requiring 7.5 Go using an Intel Core i7-7500U CPU cadenced at 2.7 Ghz with 16 Go RAM configuration.

### Microfluidic chip fabrication

The mold was fabricated on an aluminium subtrate using a computer numerical control machine (MECANUMERIC, Charlyrobot DMC300, Marssac-sur-Tarn, France) to obtain 400 μm of height. A 2 μm layer of Parylene type C was deposited31 using an evaporator (SCS LABCOTER, Indianapolis, PDS 2010, USA) to suppress milling surface defects and to favour PDMS (SYLGARD®, 184 Silicon Elastomer Kit, Dow Corning, Midland, USA) pilling. PDMS was fabricated using a standard process with a 10:1 elastomer base-curing agent ratio (w/w). The mixture was degassed and poured onto the mold up to approximately 5 mm thickness. The sample was cured at 95 °C for 1 h. The PDMS channel layer was punched (1.5 mm outer diameter, KAI Medical, Japan) to create media input and output ports (Inlet and Outlets respectively), cleaned with isopropanol and dried overnight. The PDMS was subsequently activated with air plasma and bound to a glass microscope slide to seal the channels. The chips were equipped with silicon tubing 0.5 ID (IBIDI, 10840, Gräfelfing, Germany) and proper sealing was confirmed after perfusion with water.

### Microfluidic network

Fig. 2 gives a schematic representation of the experimental set-up (see Supplementary Materials for pictures of the full set-up, Fig. S1). This network is made up of a Falcon tube (15 mL) with a microfluidic adapter (CAP) (DARWIN microfluidics, LVF-KPT-S-4, Paris, France) equipped with two plugs 1/4-28 (DARWIN microfluidics, CIL-P-309, Paris, France). These two plugs connect the inlet (IT) and outlet (OT) tubings to the remaining of the network. Both PTFE 1/32 ID tubings (DARWIN microfluidics, LVF-KTU-15, Paris, France) are equipped on both ends with 1/4-28 fitting associated with ferrules 1/16 OD (DARWIN microfluidics, LVF-KFI-06, Paris, France) (FA). IT is 45 cm long while OT is 50 cm long. Importantly, to prevent flushed worms being recaptured into the system, OT is totally immersed down to the very bottom of (~10 cm) the medium reservoir whereas IT is immersed by only 7.5 cm. IT is connected through FA to a shut-off valve (DARWIN microfluidics, LVF-KMM-08, Paris, France) (Valve). Worms are introduced into the circuit through a connector (Worm stock) composed of a Luer Lock Male connector (IBIDI, 10826, Gräfelfing, Germany) (LLM) and a Luer Lock Female adapter 1/4-28 (DARWIN microfluidics, CIL-P-618, Paris, France) (LLF). Silicone tubing 0.5 ID (IBIDI, 10841, Gräfelfing, Germany) connects the worm stock connector to the chip inlet as well as the chip outlet to the peristaltic pump (ISMATEC, IPC-N8 ISM936 Wertheim, Germany). A fitting 6-40 to 1/4-28 (DARWIN microfluidics, LVF-KFI-08, Paris, France) (FS) associated with FA connects the silicon tube to a PTFE tube (40 cm long). Connection to the pump is obtained through the association of FA with a 1/4-28 union (DARWIN microfluidics, LVF-KFI-10, Paris, France) (FU) and a 1/4-28 to 3/32 fitting (DARWIN microfluidics, LVF-KFI-06, Paris, France) (FB). The peristaltic pump tubing is 1.6 mm ID (ISMATEC, SC0425, Wertheim, Germany). Connection of the outlet of the pump to the PTFE 1/32 ID tubing OT is made with a FB+FU+FA association.

**Fig. 2:**
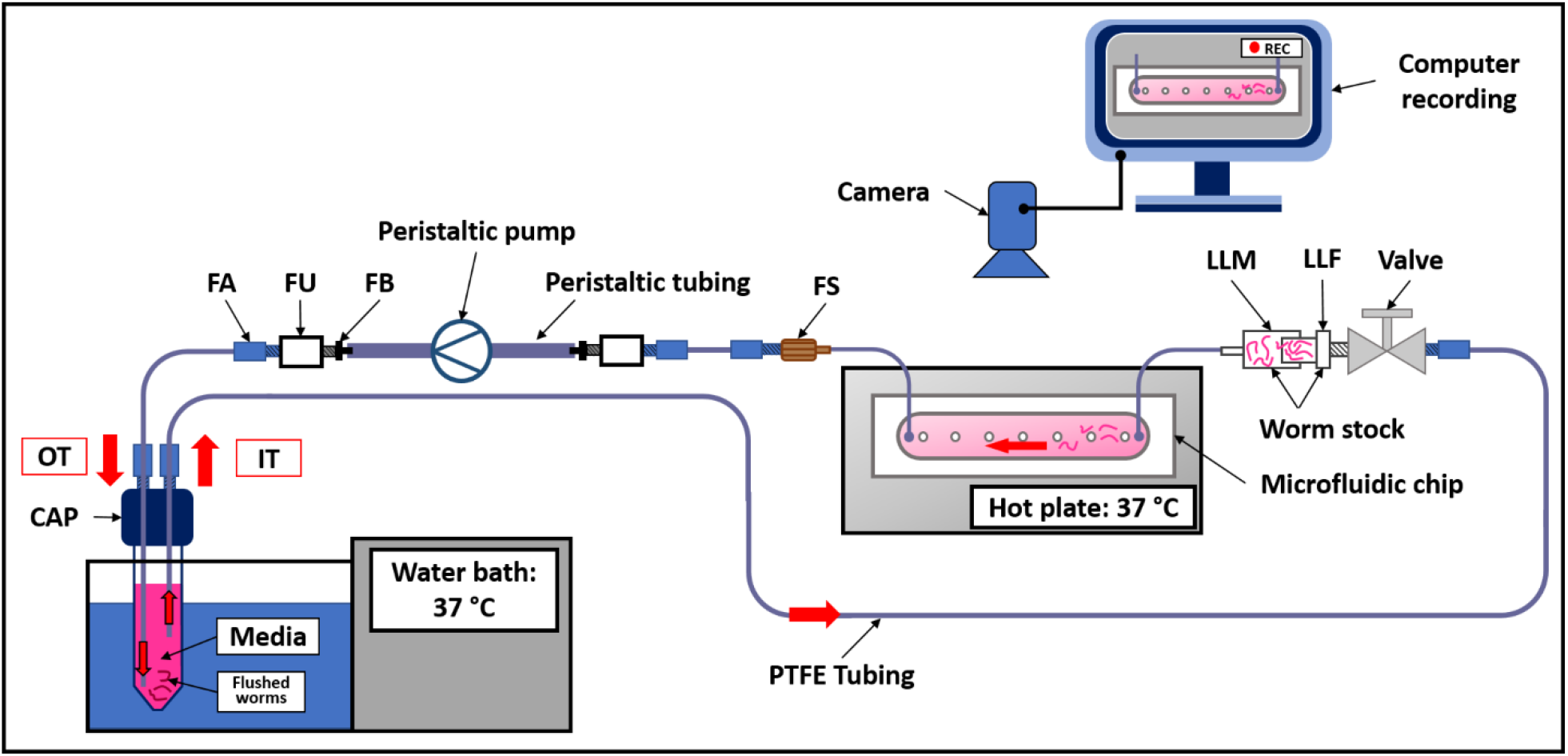
Schematic representation of the microfluidic experimental set-up. The culture medium supplemented or not with PZQ or ART was maintained at 37 °C using a water bath and pulled with a peristaltic pump into the microfluidic chip at 1 mL.min-1. Chips were placed on a heating plate. The detachment of the worms in the microfluidic chip was video-recorded for 1 hour. Components of the microfluidic network are described in Fig. S1.

### Device surface coating

Chip chambers were coated with 250 μg.mL^−1^ of Type I-A collagen (FUJIFILM, Cellmatrix Type I-A, 631-00651, Neuss, Germany) solubilized in aqueous acetic acid (0.02 M) and introduced with a 1 mL syringe into the chip. After incubating at room temperature for 3 h, the chips were rinsed twice with 3 mL of water. Chips were then connected to tubings and primed with 10 mL of complete RPMI medium prior to the connection of IT to CAP. For chips coated with MDCK cells, the inlet port was disconnected from tubing to introduce the MDCK cells (10.000 cells/chip) with a 1 mL syringe, then reconnected to the tubing. Finally, the circuit was connected to a peristaltic pump (ISMATEC, IPC-N8 ISM936 Wertheim, Germany) without flow for 3 h. Unattached cells were then flushed at 35 μL.min^−1^ for 48 h, after which the 15 mL stock tube was replaced with a new one containing 9 mL of dedicated medium.

### Worm introduction into device

After having conditionned the microfluidic system, the peristaltic pump was stopped. The valve was closed and the circuit opened at the worm stock connector (disconnection between LLM and LLF). A 1 mL pipette tip was shortened approximately by 0.5 cm, and used with a pipette to collect 30 couples and introduce them into LLM. To prevent bubble introduction into the circuit, LLM and LLF were filled by pipetting media before reconnection. The valve was opened again and the microfluidic chip connected to the peristaltic pump (ISMATEC, IPC-N8 ISM936 Wertheim, Germany). The pump was set for pulling the culture media at a flow rate of 75 μL.min^−1^ and placed with the chip in an incubator to reduce tubing length and temperature drift (37 °C, humid atmosphere, 5% CO_2_). The system was primed for 30 min before starting the worm adhesion assay. Importantly, to avoid worms clogging the tubbing and to disperse the worms within the chip, the system was placed vertically to let the couples settle by gravity. For that, the worm stock and microfluidic output were placed upwards and the microfluidic input placed downwards as shown in Fig. S2.

### Adhesion studies on different surfaces

For each experiment, 30 couples were introduced into the chips (n=2) with different glass surface coatings: w/o coating, coated with MDCK cells, with type I-A collagen or with both MDCK cells and type I-A collagen. Chips were placed on a heating block set at 37 °C (Major Science, MD-02D, USA), and connected to 15 mL tubes placed in a water bath set at 37 °C as depicted in Fig. 2. A camera (Microsoft, Q2F-00015, USA) was used to monitor the microfluidic channel at the beginning of the experiment in order to identify worms that lost adherence and to measure the duration of their residence in the microsystem before being flushed out by the flow. To modulate flow conditions, the peristaltic pump was programmed to pull the culture medium at different flow rates: 1) 50 μL.min^−1^ to 300 μL.min^−1^ with a step of 50 μL.min^−1^ every min; 2) 300 μL.min^−1^ to 1 mL.min^−1^ with a step of 100 μL.min^−1^ every 2 min; 3) 1 mL.min^−1^ to 3.4 mL.min^−1^ with a step of 200 μL.min^−1^ every 3 min. Percentages, mean values and standard deviations were calculated using RStudio (1.1.463 release, RStudio Inc., Boston, USA). The percentages of worms still attached to the chip were plotted against the flow rate. A 2 segment linear regression analysis was performed using SigmaPlot v14.5 with Piecewise fitting curve option to identify the transition value (T1, see supplementary table 1). We next performed a linear regression using the data between 0 and 1.6 mL.min^−1^ for glass, MDCK and collagen + MDCK, while the data between 500 μL.min^−1^ and 1.6 mL.min^−1^ values were used for collagen. This analysis furnished the slope of the first segments, the standard deviation and the 95% confidence intervals (see supplementary table 2).

### Drug testing under flow conditions

Praziquantel (PZQ) (SIGMA, P-4668, USA) was prepared as a 0.5 M stock solution in DMSO and diluted in RPMI Horse serum from 900 nM to 25 nM. Artemisinin (ART) (Tokyo chemical industry, A2118, Japan) was prepared as a 0.1 M stock solution in DMSO and diluted from 900 μM to 12.5 μM in RPMI Horse serum. Stock solutions were stored at 4 °C. Once the worms were introduced in the system as previously described, the flow rate was increased up to 1 mL.min^−1^ by 200 μL.min^−1^ increments every 5 s in order to favor worm attachment to the collagen coated chips. The recording was started and the 1 mL.min^−1^ flow was maintained for 5 min. This lag period allowed the elimination of weakened worms. Next, PZQ or ART were added to the tube containing the culture medium. The medium was mixed and infused into the device. The video recordings were used to count the worm couples present in the microfluidic chamber as a function of time. Percentages, mean values and standard deviations were calculated using RStudio (1.1.463 release, RStudio Inc., Boston, USA). Experiments were performed in triplicate.

### Drug testing under static conditions

Typically, 10 worm couples were incubated in 12 well plates at 37 °C for 120 h in RPMI Horse supplemented with PZQ or ART (0 to 900 nM for PZQ or 0 to 900 μM for ART, 3 mL per well). The media were removed and renewed every 24 h. After this step, each well was immediately recorded for 15 s under a macroscope (OPTIKA, SZ6745TR, Italy) equipped with a camera (Microsoft, Q2F-00015, USA). In this experiment, we used a more resolved ‘severity score’ than those classically used for drug evaluation against helminths in static conditions^27^. We designed this 5 level scoring since it gives a more precise evaluation of phenotypes, in particular paralysis or tegument alterations (S4 = Normal mobility and blood sucker adhesion; S3 = reduction of mobility and/or loss of blood sucker adhesion; S2 = minimal mobility or occasional movements; S1 = no mobility except intestinal movement; S0 = total loss of mobility, no movement, death). RStudio (1.1.463 release, RStudio Inc., Boston, USA). was used for calculating percentages, mean values and standard deviations.

## Results

### Chip design

In order to generate a chip with a dynamic flow that forces worms to attach to the walls, we chose a simple design based on a single channel that can host a sufficient number of worms (30 to 40 couples) to improve the statistical significance of analysis. The chip had to be small enough to accommodate regular cameras and microscopes and to be parallelized. Thus, we designed a 1 cm × 5.8 cm single channel supported by 7 pillars to prevent any sagging of the chip (Fig. 3a). In order to ease the elimination of the worms by the flow stream during treatments, we chose a 400 μm height for the channel that allows the circulation of damaged and/or swelling worms. This chip design enabled also a quick, safe and easy introduction of the worm couples that reduces manipulation as much as possible, and thus the risk of worm damage.

**Fig. 3:**
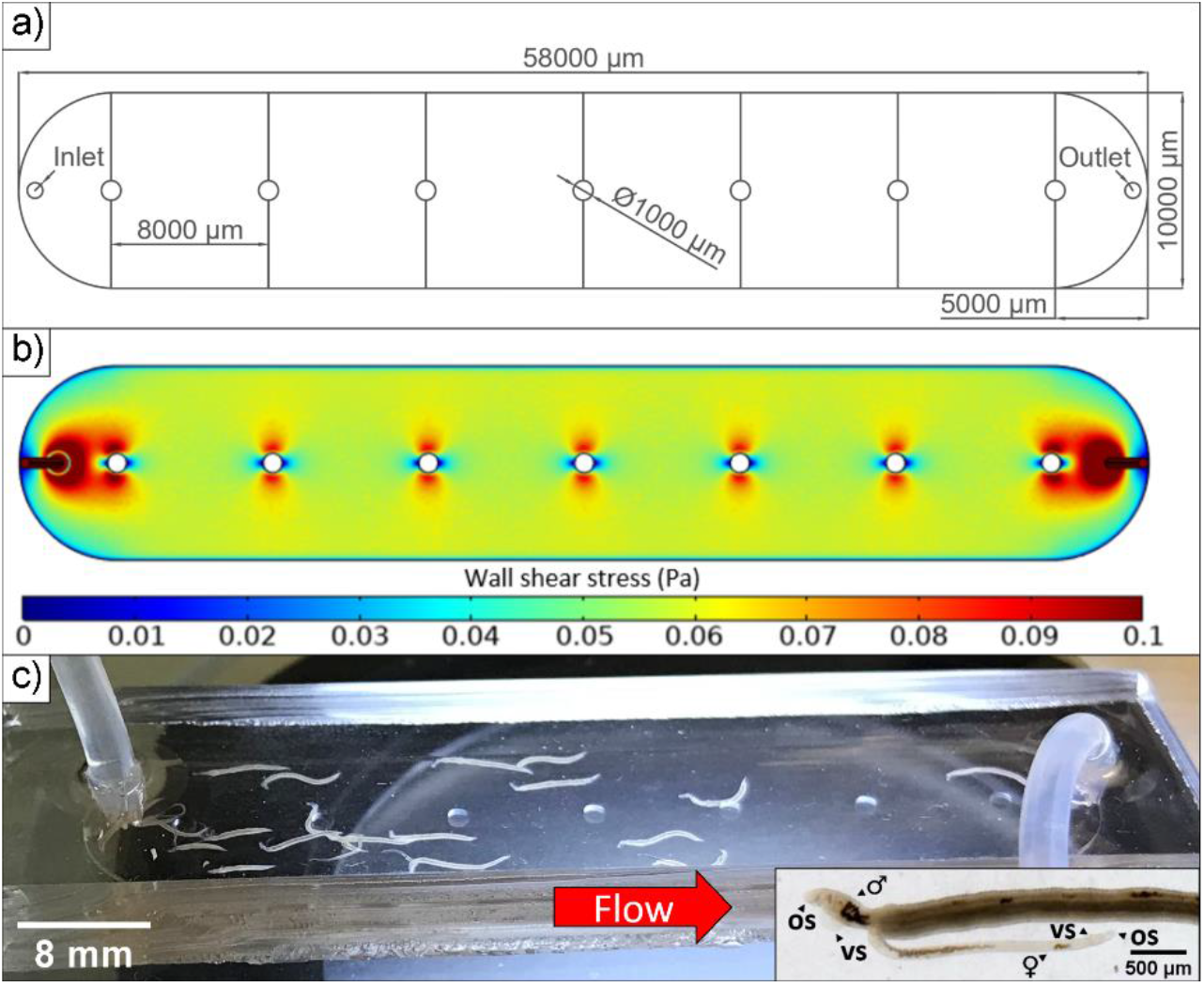
Microfluidic chip model. a) Layout of the microfluidic device. Chamber of 58.000 μm of length, 10.000 μm of width, 400 μm of height and a pillar of 1000 μm of diameter every 8000 μm. Media input (Inlet) and output (Outlet) ports location; b) Numerical simulation of the wall shear stress (WSS, colour gradient in Pa); c) Couples of schistosomes under flow conditions loaded in a collagen coated chip (1 mL.min-1). Insert: Males 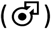 and females 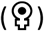 attached to chip surface using their oral (OS) and ventral suckers (VS).

The flow rates generated in our system were simulated using a volumic mass of 1007 kg.m^−3^ and a dynamic viscosity of 0.958 mPa.s for the culture medium^32^. Supplementary table S3 gives the values of the wall shear stress WSS (Pa), maximum velocity (m.s^−1^), Reynolds number and pressure gradient (Pa.m^−1^) obtained for different flow rates. The flow was found to be laminar (low Reynolds number) and the range of WSS ranged between 0.003 and 0.206 Pa (Fig. 3b and Fig. S6). Thus, the range of flow rates used in our system reproduce the WSS conditions encountered by worms in the human blood system and in particular in veins and venules^33^.

Finally, we showed that under dynamic conditions, fresh worms firmly attach to the walls, with a strong preference for the glass surface, and actively resist the flow (Fig. 3c). It worth noting that most of worm couples stayed paired, which is a sign of health, and that both males and females contributed to the attachment through their suckers. Immediately after starting and increasing the flow, damaged and/or weak worms were eliminated in few minutes from the chip (10 to 15%) and were flushed out and pushed inside the outlet tubing, whereas heathy worms remained firmly gripped to the chip walls for hours.

### Optimization of worm storage in *ex-vivo* conditions

One of the major limiting factors to performing studies on adult schistosomes is their short survival period in culture media after perfusion from their vertebrate host. Since a continuous perfusion of “fresh worms” is not ethically or practically feasible, proper storage of a sufficient worm population for several days is a critical element to compare successive experiments. Consequently, the culture of worms in an optimal medium is a key point for reliable experiments. Although several studies have aimed at identifying positive and negative factors contained in various sera^34–36^ for worm culture, no firm conclusions regarding the most appropriate serum to use in our experiments could be drawn from previous reports. Therefore, and before starting the microfluidic experiments, we characterized different media to identify those enabling the maintenance the adult worms in a healthy status for at least several days after recovery from infected hamsters. We varied the cell culture basal medium (M199, DMEM and RPMI) and the serum additive (10% of heat-inactivated fetal calf, horse or human serum). Figure 4 shows the worm viability as a function of time for the different serum additives added to RPMI (see Fig. S3 for M199 and DMEM). The data show that the serum type has a major impact on worm longevity. Worms maintained with human serum showed a survival rate greater than 90% whatever the medium composition. Horse serum achieved a similar performance except when combined with M199 medium which gave less than 90% of survival after 25 days of culture. Beyond this period, the viability of couples dropped drastically for all the culture media tested with, however, a better result for the RPMI and DMEM media. Note that calf serum, which is classically used for schistosome cultures, showed the poorest results with a survival rate of 70% at 29 days with RPMI and less for the other culture media tested. Consequently, since our adult worms are obtained on a weekly schedule, we selected the combination of RPMI medium and horse serum for subsequent experiments. Note that horse serum has the advantage of being easily accessible and safer to use than human serum.

**Fig. 4:**
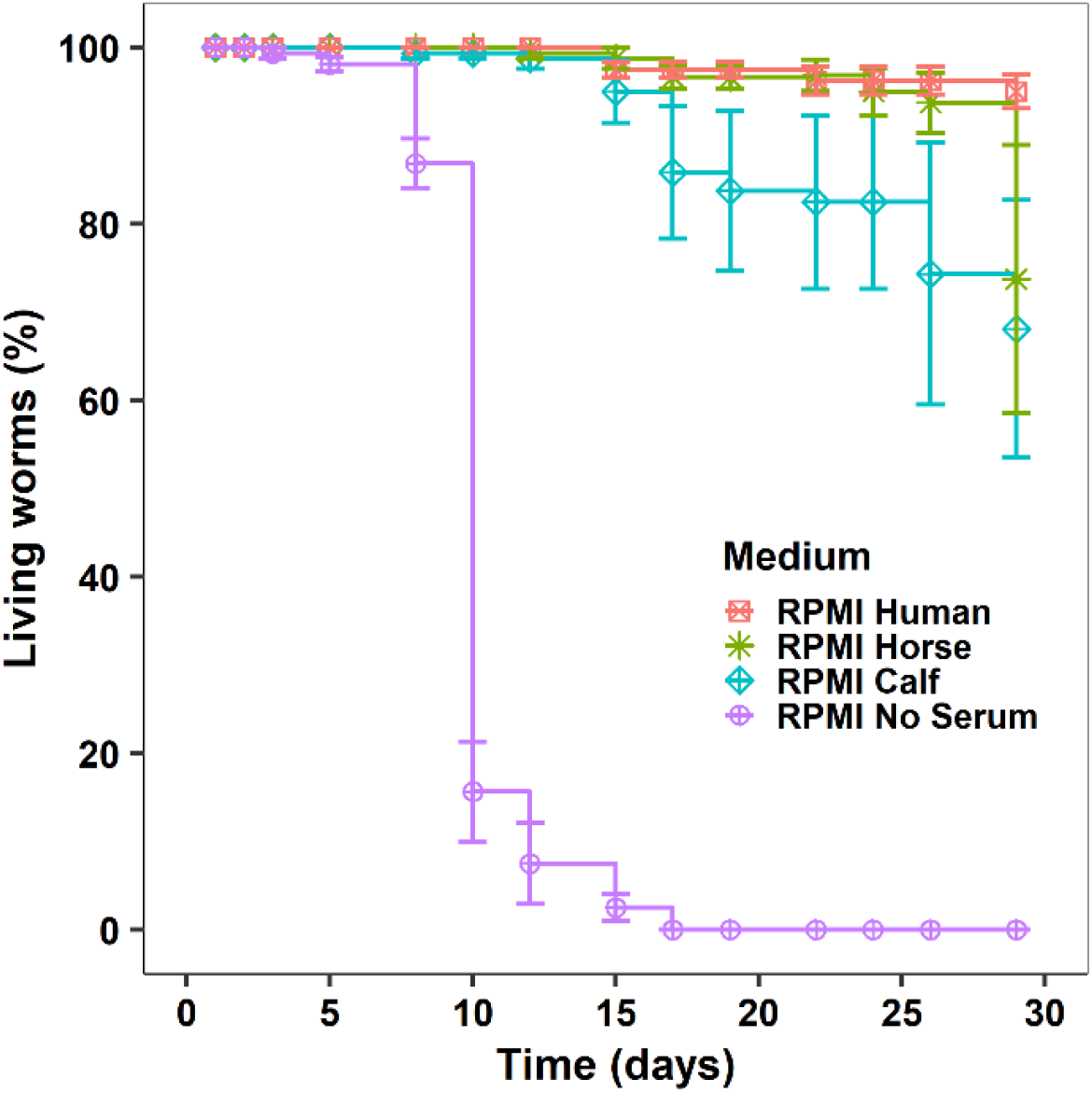
Progression of the survival rate of S. mansoni pairs as a function of time for RPMI culture media supplemented or not with serum in static Petri dish culture conditions.

### Worm adhesion capabilities in microfluidic chips

The ability of the worm couples to grip the walls of the microfluidic system potentially depends on its geometry but also on the physicochemical nature of the walls constituting the perfusion channel. Indeed, *in vivo*, adult worms mostly attach to the vein endothelium using their oral and ventral suckers together with a peristaltic action of the tegument^28^. In our system, without a confined geometry like that of veins, we logically observed that the attachment mostly relied the ability of suckers to grip the chip surface (See supplementary movies S1 and S2). This led us to examine the influence of different surface coatings on the resistance of the worms to the flow through the measurement of their residence time in the device.

Fig. 5 gives the number of worm pairs in the chip as a function of the flow rate for the different coatings (raw glass, collagen, MDCK cells or MDCK cells on collagen). The responses of the worms to the flow showed a biphasic behaviour. A first linear response with a negative slope was immediately observed for the raw glass, glass+MDCK and collagen+MDCK coated chips, with a transition value of the slopes occuring at ~1.6 mL.min^−1^. (See Figs. S4, S5 and table S2). In contrast, the collagen coated chip was characterized by an initial plateau at 100% residence of worm pairs from 0 to 0.4 mL.min^−1^ flow rate, followed by a linear response to the flow with a negative slope. The slope of the segment is a quantification of the resistance of the worms to the flow. For glass and glass+MDCK coated surface, the loss of worms (56 and 57 % loss per mL.min^−1^) is much larger than those occuring with the collagen and collagen+MDCK surfaces (38 and 22 % loss per mL.min^−1^). The initial plateau for the collagen chip suggests an easier initial gripping of the worms when the flow starts and increases. We tried to understand why such a difference exists in the worm gripping between the uncoated and the collagen coated chips, independantly of a cell layer. Close examination of the chips under the microscope revealed that worms firmly gripped the cell layer. However, after starting the flow, on the chips not coated with collagen, the cell layer “pilled off” under sucker traction and worms were unable to reattach. Surpringingly, this phenomenon of cell detachment upon sucker traction was also observed on the collagen+MDCK coated chips, making the cell layer heterogeneous and likely explaining the higher standard deviation in the data compared to the simple collagen coating chips. Therefore and counterintuitively, these data show that a simple collagen coating on a glass surface is sufficient to promote robust worm gripping by their suckers whereas a cellular coating does not favour robust gripping and rather adds complexity with no significant advantage.

**Fig. 5:**
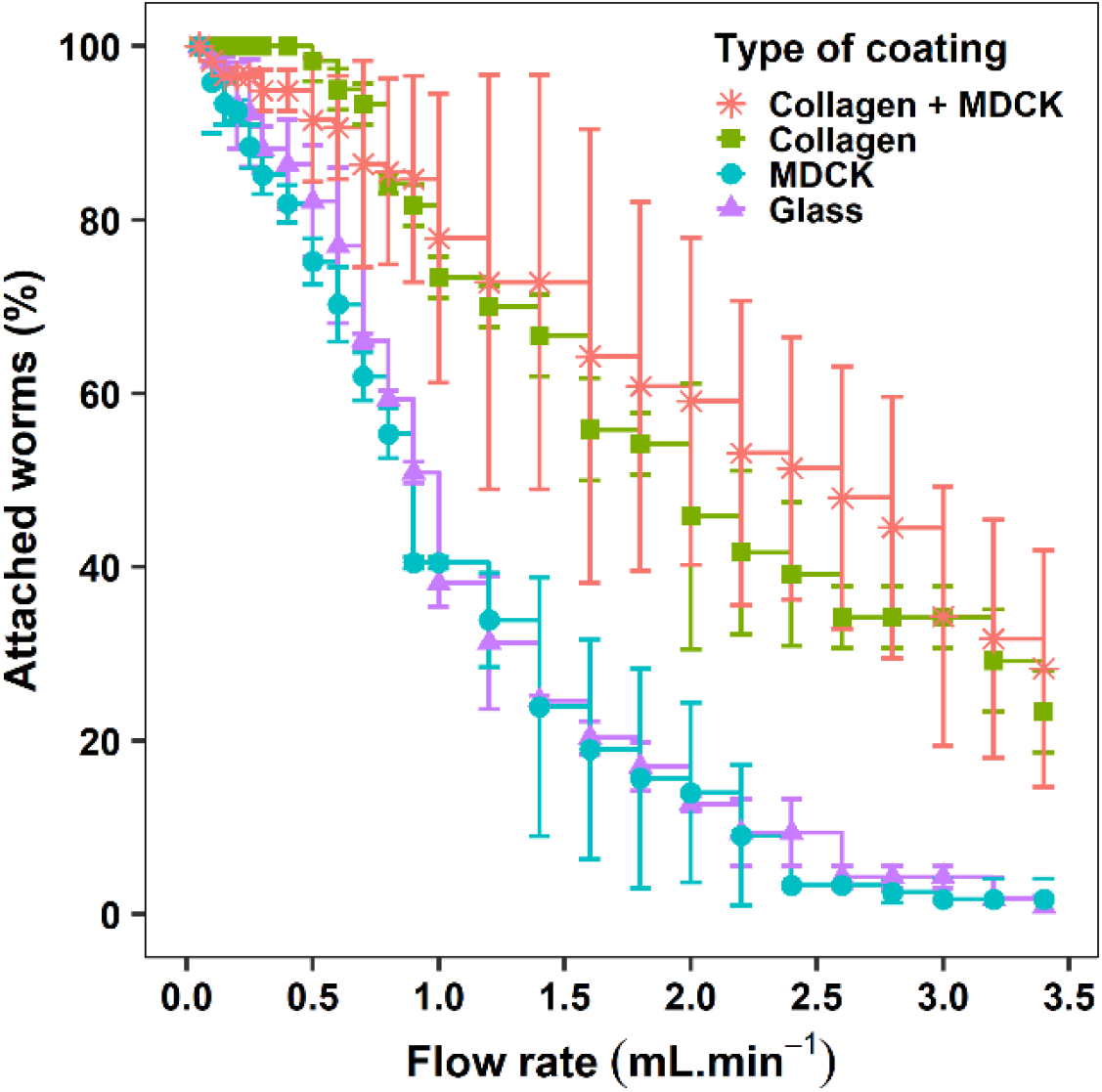
Evolution of the number of schistosome couples in the microfluidic chip as a function of flow rate for different internal wall coatings: None, collagen, MDCK cells and MDCK cells + collagen.

For the rest of this study, we selected a collagen surfacing which is easier to implement and gives robust and reproducible worm gripping at a flow rate of 1 mL.min^−1^. This flow rate allowed a >80% worm retention but still forced firm gripping from the worms and allowed elimination of weak worms within a few minutes.

### Determination of early effects by drugs in microfluidic device

Having defined the most reliable surface coating and flow rate for promoting worm residence in the microfluidic channels, we measured the effect of two well known therapeutic molecules, PZQ and ART, on the ability of worms to grip to the chip surface under dynamic conditions. For these experiments, the microchannels were infused at a flow rate of 1 mL.min^−1^ and the number of worms remaining in the microchannel was measured as a function of time for different doses of PZQ or ART (Fig. 6a, b f and g).

**Fig. 6:**
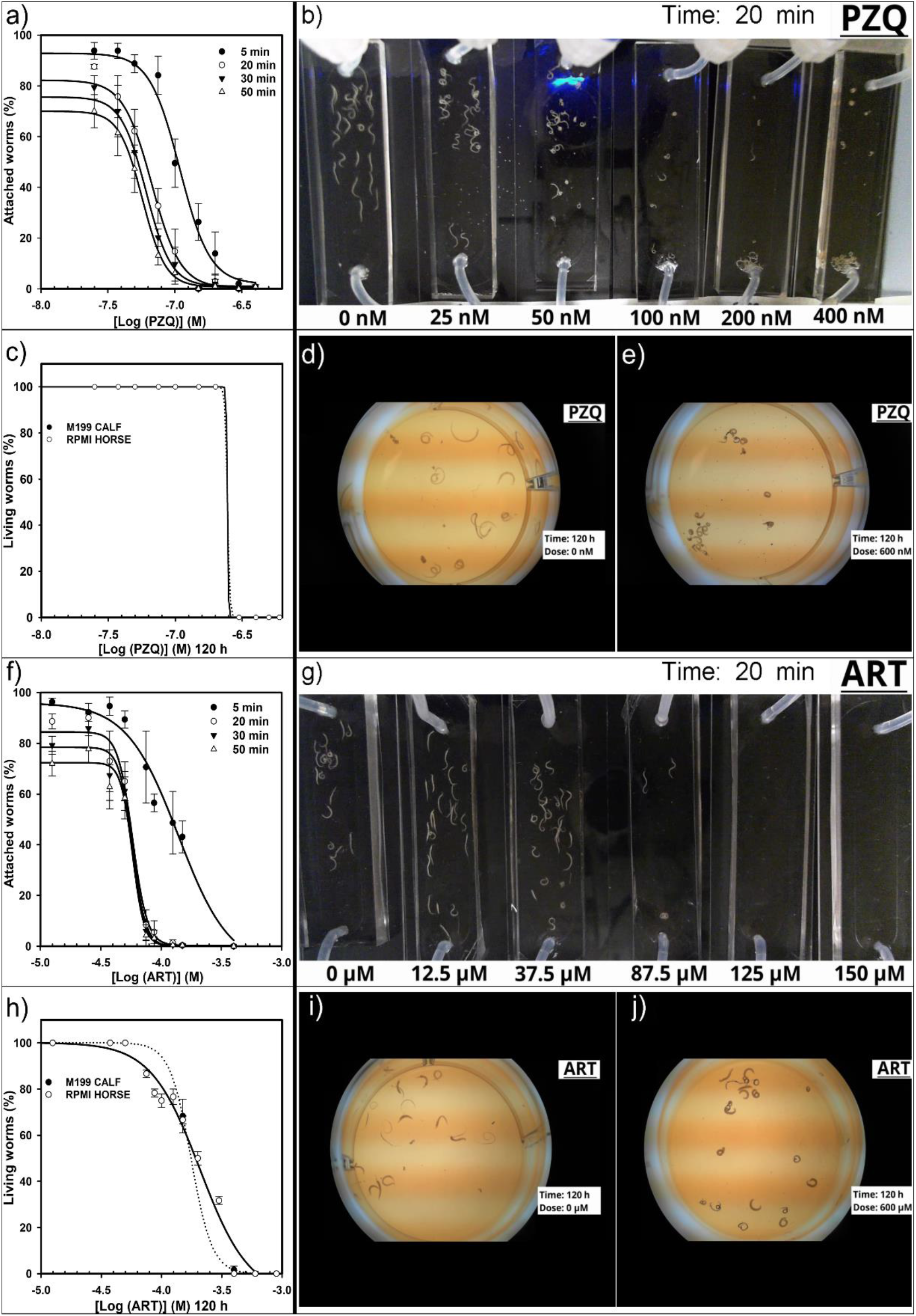
Microfluidic and static IC50 determinations of responses to PZQ and ART. a) Microfluidic determination of IC50 based on percentage of schistosomes attached to collagen chip surface at 5, 20, 30 and 50 min depending on PZQ doses; b) Representative image of 6 chip series at 20 minutes using six doses of PZQ (0, 25, 50, 100, 200, 400 nM) with 30 couples per chip at 1 mL.min-1 flow rate; c) Static determination of IC50 based on percentage of living schistosomes at 120 h depending on PZQ doses; d) Representative image of 1 well of a PZQ negative control at 120 h with 10 couples; e) Representative image of 1 well with a 600 nM PZQ dose at 120 h with 10 worm couples; f) Microfluidic determination of IC50 based on percentage of schistosomes attached to collagen chip surface at 5, 20, 30 and 50 min depending on ART doses; g) Representative image of 6 chip series at 20 minutes using six doses of ART (0, 25, 50, 100, 200, 400 nM) with 30 couples per chip at 1 mL.min-1 flow rate; g) Static determination of IC50 based on percentage of schistosomes at 120 h depending on ART doses; i) Representative image of 1 well of an ART negative control at 120 h with 10 couples; j) Representative image of 1 well with a 600 nM ART dose at 120 h with 10 couples.

We observed that a PZQ dose of 25 nM had no significant effect on worm gripping compared to the control (approximately 8% of the initial worms number were flushed out after 5 min and 30% after 50 min). The effect of PZQ started to be observed at 37.5 nM with 60% of the worms remaining in the system after 50 min (Supplementary movie S3). At 50 nM, we began to perceive erratic contractions of the worms and 50% of the worms were able resist the flow after 50 min. From 100 to 400 nM, a rapid paralysis and thus detachment of the worms was observed, together with a significant swelling and a reduction in their length. Consequently, only 20% of the worms still remained after 10 min, while they were all carried away by the flow after 50 min. The IC_50_ values for 20, 30 and 50 min residence times are reported in Table 1. Importantly, we noticed that after 20 min of treatment, IC_50_ values were similar and close to 60 nM, suggesting that the effect of the drug on the worms is optimal after 20 min of perfusion but this can be extended if needed.

**Table 1:** Residence IC50 values determined in dynamic conditions. These values correspond to the concentration of PZQ or ART that induced 50% of worm detachment from the collagen coated chips after 5, 20, 30 or 50 min of treatment at a flow rate of 1 mL.min^−1^ in RPMI horse medium. Data are presented with their 95 % confidence intervals (CI).

|  | PZQ | ART |
| --- | --- | --- |
| Time (min) | IC<sub>50</sub> ± 95% CI (nM) | IC<sub>50</sub> ± 95% CI (μM) |
| 5 | 109 ± 13 | 139 ± 42 |
| 20 | 67 ± 10 | 58 ± 5 |
| 30 | 61 ± 9 | 58 ± 5 |
| 50 | 57 ± 7 | 58 ± 6 |

Next, we evaluated the effect of ART on worm gripping under microfluidic conditions. Worms started to detach from the chip at a dose of 50 μM (Fig. 6f and g). At this concentration, fewer than 50% of the worms remained after 50 min (Supplementary movie S8). Above 75 μM, the effect of the drug on worm morphology began to be noticeable and led to a massive flush out (>90%). Again, the calculated IC_50_ values were almost identical over time at 58 μM after 20 min of chip perfusion.

In parallel and using the same batch of worms, we also determined IC_50_ values for PZQ or ART using the classical method in Petri dishes under static conditions. We observed a very sharp effect of PZQ since after 120 h a dramatic change in the scoring occurred between 150 nM and 300 nM (Supplementary movies S4-S7), leading to an IC_50_ value of 245 nM.

The effect of ART on worm viability was much more gradual. Worms started to degrade and to die from 12.5 μM up to 400 μM where all worms were classified as dead (Supplementary movies S9-S12). This resulted in an IC_50_ value of 200 μM after 120 h of exposure to ART. Since these experiments in static conditions lasted 120 h and to ensure that our IC_50_ measurements were not influenced by medium composition, we also determined the IC_50_ in M199 with calf serum. The values obtained were almost identical and the corresponding IC_50_ values for PZQ and ART from these Petri dish experiments are collected in Table 2.

**Table 2:** Worm death IC_50_ values determined in static conditions. These values correspond to the concentration of PZQ or ART that induced 50% of worm death using a 5 level severity-score in 6 well Petri dish format after 120 h of treatment in RPMI horse and M199 calf medium. Data are presented with their 95 % confidence intervals (CI).

|  | PZQ (120 h) | ART (120 h) |
| --- | --- | --- |
| Medium/serum | IC<sub>50</sub> ± 95% CI (nM) | IC<sub>50</sub> ± 95% CI (μM) |
| RPMI/horse | 245 ± 5 | 176 ± 30 |
| M199/calf | 245 ± 1 | 208 ± 24 |

Even if the measured phenotypes (attachment vs survival) are different and thus difficult to compare side by side, the IC_50_ values obtained in dynamic conditions are in the same range as those obtained in ‘classical’ static condition for PZQ and ART. This suggests that our microfluidic device allows anticipation of drug effects in less than 20 min.

## Discussion

The identification of new drugs to treat schistosomiasis is one of the priorities of WHO for parasite control37. Despite decades of research to identify an alternative to PZQ, this molecule remains the only one recommended to treat this parasitosis38. The mode of action of PZQ has been recently elucidated,39 more than 40 years after the observation of its antihelminthic activity40. PZQ acts on the transient receptor potential channel (TRPM), whose activation instantaneously paralyses worms and leads to death by altering their surface tegument and giving access to the immune system. Therefore, the design of PZQ derivatives, possibly more potent and specific for the schistosome target, are ongoing41. However, the identification of new drugs remains highly limited by the absence of simple and practical screening/analysis methods dedicated to adult schistosomes42. Drug selection still remains dependent on the observation of complex phenotypes and/or sophisticated image analysis of worm motility, egg laying or pairing18. In addition, the short viability of adult worms in in vitro culture systems and the rapid loss of their biological functions strongly biases the results, particularly in the case of slow acting compounds. Selection and functional characterization of active coumpounds on whole worms are thus a challenge.

Monitoring the state of a living cell assembly at the level of a single cell, tissue, organ or whole body is carried out by quantifying various parameters usually identified by fluorescent or radiolabeled markers. These techniques are often destructive for the sample and performed at the end of the experiment. Label-free and real-time techniques have developed significantly for several years43 and, in particular, applied in the field of organ-on-a-chip27. Impedance spectroscopy (IS) is such a label-free technique suitable for quantifying the properties of cell assemblies in real time30. It is compatible with high-throughput analyses and allows kinetics of the biological process to be measured and studied.

However, given the complexity of a cell assembly in a living worm, the interpretation of electrical data and the link to biological processes remain difficult to disentangle in many configurations. These methods are under development for monitoring schistosomula viability.44 Their parallelization should provide an effective tool for the screening of drug libraries43,44. Despite these new technological advances, application to older worm stages (juvenile and adults) and particularly paired couples is very unlikely due to their large size (~1 cm).

The study of whole animals in a smart microfluidic system is a very recent research topic in the field of systems biology. Some basic systems have been reported for the study of the free-living Caenorhabditis elegans worm.45–46 In contrast, studies on trematodes are scare and mostly focused on worm locomotion mechanistics.28 In parallel, progress has been made on the use of microfluidics for the detection of the parasites in human fluid either by microfluidic PCR, antigen identification or egg isolation47. The application of microfluidic technologies to the study of adult worm couple viability and fertility, or the evaluation of drug effects remains to be explored.

One significant limitation for adapting these microsystems to the study of adult worms is the limited viability of adult worms “ex-vivo”. Therefore, the analysis of long lasting effects of drugs can be difficult to distinguish from the natural degradation of worms outside their host. Culture media have been optimized to allow the in vitro maintenance of adult worms for a few weeks, using worm mobility together with a sustained egg production as the main criteria. However, none of these optimizations led to the production of viable eggs suggesting that medium optimization is not yet sufficient for long term maintenance of parasites ex vivo27. Therefore, we have chosen to design a fluidic system that allows a quick measurement of a basic phenotypic parameter: the surface attachment by suckers. This system necessarily selects healthy worms (males, females or couples) since it promotes the worm muscular reflex to attach onto the surface in a dynamic environment mimicking the blood flows occurring in vivo. In the portal system and in particular in the mesenteric vein where parasites resides, the flow speed is about 0.001 m.s-1 and WSS estimated to be around 0.3 Pa48,33. Reproduction of these flow conditions is absolutely necessary to ensure that the worms are under realistic mechanical stress conditions that make the test more predictive and avoid over- or underestimation of the effect of molecules on the adhesion capacity of worms. In particular, we showed that a simple collagen coating strongly improves the ability of worms to firmly grip with their suckers onto the glass surface even at high flow rates. While this parameter cannot be considered in static systems, it is an easy and powerful indicator of worm health and viability that has been overlooked. A major technical advantage of our system is that it does not require sophisticated or time-consuming image analysis since scoring is based on a binary method of evaluation: attached or not.

We chose to validate our system using the two major drugs used against the parasite, PZQ and ART. These compounds have different modes of action. PZQ triggers a sudden calcium efflux that paralyses worms and thus has an immediate effect on worm attachment capability. In contrast, the mode of action of ART is more pleiotropic with the production of free-radicals that requires the presence of heme in the worm gut, a byproduct of hemoglobin degradation49. The production of free-radicals induces both oxidative and metabolic stresses and results in damage to tegument, tissues and reproductive organs 50–51. Thus contrary to PZQ, ART was not expected to promote an immediate effect on worms. Surprisingly, we observed a quick effect on worm attachment in a time and concentration dependent manner for both compounds. Moreover, we obtained IC50 values under flow conditions that are significantly lower than those measured under static conditions. For example, we obtained for PZQ an IC50 of 57±7 nM at 50 min with the fluidic system compared to an IC50 of 245±1 nM in the static system at 120 h. Similarly, ART gives an IC50 of 58±6 μM in the device compared to an IC50 of 176±30 μM in a Petri dish at 120h in M199 calf serum. These results highlight the difficulty in comparing phenotypic assays performed under different conditions. Our microfluidic system must be considered as a novel phenotypic assay that is particularly sensitive due to the measurement of worm attachment capability.

One asset of the system described here is that it allows parallelization and simultaneous comparisons in using a larger number of live worms (30 to 40 couples) compared to Petri dish assays (5-10 couples), facilitating the reproducibility of experiments.12

Evidently, our dynamic system does not prejudge the delayed lethal effect of tested compounds but rather gives a direct insight into their toxicity and visualizes an immediate effect compared to long term experiments in Petri dishes. Besides the sensitivity of worm gripping to drugs, other factors can contribute to the differences observed between flow and static conditions. Indeed, the flowrate has probably a major effect on drug distribution and metabolization compared to static conditions. Schistosomula and adult schistosomes have a very active metabolism characterized by a massive glucose consumption52 associated with lactate production53 and excretion54. However, high lactate production and its accumulation55 due to a low diffusion rate in a static environment can generate local acidification around worms. This acidification is well described to modify metabolization of drugs in cancer treatments56. Under flow conditions, the permanent actuation of drugs and culture medium probably reflects more the physiological conditions encountered by the worms in the bloodstream.

Finally and conveniently, flushed out worms can be retrieved from the devices and further investigations such as metabolic or image analysis can be performed. In addition, exchange of active compounds or pulse-chase treatment sequences can be easily performed in such a device. Finally, the entire device (chip and tubing) consumes a very small amount of circulating fluid (5-10 mL) and thus reduces the costs associated with the use of elaborate media and/or active molecules.

Having established the proof of concept that our dynamic microsystem is reliable for the identification of active molecules against the parasite, we now aim to test further series of drugs together with the implementation of automated worm counting by simple real time image analysis.

## Conclusions

Selection and characterization of schistosomicidal drugs still remain major challenges mostly due to the limitations of relevant screening systems together with the poor longevity of the parasite in vitro. Classical screenings in Petri dishes are cumbersome since they rely on the optical analysis of complex phenotypes mostly based on worm motion and/or tissue integrity. Therefore, the design of novel systems enabling a practical and rapid identification of molecules acting on the parasite is desirable. We report such a system that can be parallelized with a large number of worms and relying on a simple phenotype, i.e. the attachment of worms to the chip surface under flow conditions. It is also affordable, and thus should facilitate the identification of new drugs against this re-emerging parasite.

## References

1 https://www.who.int/news-room/fact-sheets/detail/schistosomiasis

2 D.G. Colley, et al., Lancet, 2014, 383, 2253–2264.

3 J. Boissier et al., Lancet Infect Dis., 2016, 16(8), 971–979.

4 C.S. Nation, et al., PLoS Negl. Trop. Dis., 2020, 14(4), e0007951.

5 A.J. Molehin, J. BioMed. Sci., 2020, 27:1–7.

6 P. Mäder, G.A. Rennar, A.M.P. Ventura, C.G. Grevelding, M. Schlitzer, ChemMedChem, 2018, 13, 2374–2389.

7 W.E. Secor, S.P. Montgomery, Future Med. Chem., 2015, 7, 681–684.

8 W. Wang, L. Wang, Y.-S. Liang, Parasitol. Res., 2012, 111, 1871–7.

9 P.G. Fallon, M.J. Doenhoff, Am. J. Trop. Med. Hyg., 1994, 51, 83–88.

10 S.D. Melman, M.L. Steinauer, C. Cunningham, L.S. Kubatko, I.N. Mwang, N.B. Wynn NB, et al., PloS Negl. Trop. Dis., 2009, 3, e504.

11 T. Crellen, M. Walker, P.H.L. Lamberton, N.B. Kabatereine, E.M. Tukahebwa, J.A. Cotton, et al., Clin. Inf. Dis., 2016, 63, 1151–9.

12 J.T. Moreira-Filho, A. C. Silva, R.F. Dantas, B.F. Gomes, L.R. Souza Neto, et al., Front. Immunol., 2021, 12: 642383.

13 R.A. Paveley, N.R. Mansour, I. Hallyburton, L.S. Bleicher, A.E. Benn, et al., PloS Negl Trop Dis, 2012, 6, e1762.

14 S. Chen, B.M. Suzuki, J. Dohrmann, R. Singh, M.R. Arkin, C.R. Caffrey, Commun Biol, 2020, 3, 747.

15 C. Lalli, A. Guidi, N. Gennari, S. Altamura, A. Bresciani, G. Ruberti, PloS Negl Trop Dis, 2015, 9, e0003484.

16 N.R. Mansour, Q.D. Bickle, PloS Negl Trop Dis, 2010, 4, e795.

17 P.S. Ravaynia, F.C. Lombardo, S. Biendl, M.A. Dupuch, J. Keiser, A. Hierlemann, et al., Adv Biosyst, 2020, 4, 1900304.

18 B. Ramirez, Q. Bickle, F. Yousif, F. Fakorede, M-A. Mouries, S. Nwaka, Expert Opin. Drug Discov., 2007, 2, S53–S61.

19 B.J. Neves, R.F. Dantas, M.R. Senger, W.C.G. Valente, J. de M. Rezende-Neto, W.T. Chaves, et al., Med Chem Commun, 2016, 7, 1176–82.

20 G. Rinaldi, A. Loukas, P.J. Brindley, J.T. Irelan, M.J. Smout, Int J Parasitol Drugs Drug Resist, 2015, 5, 141–148.

21 T. Manneck, O. Braissant, Y. Haggenmuller, J. Keiser, J Clin Microbiol, 2011, 49, 1217–1225.

22 T.G. Geary, J.A. Sakanari, C.R. Caffrey, J. Parasitol., 2015, 101, 125–133.

23 N. Aulner, A. Danckaert, J. Ihm, D. Shum, S.L. Shorte, Trends Parasit., 2019, 35(7), 559–570.

24 J.M.F. Gardner, N.R. Mansour, A.S. Bell, H. Helmby, Q. Bickle, PLoS Negl. Trop. Dis., 2021, 15(7), e0009490.

25 F. C. Lombardo, et al., Nature Protocols, 2019, 14, 461–481.

26 G. Padalino, S. Ferla, A. Brancale, I.W. Chalmers, K.F. Hoffmann, Int. J. Parasitol. Drugs Drug. Resist., 2018, 8, 559–570.

27 J. Wang, R. Chen, J.J. Collins III, PLoS Biol., 2019, 17(5), e3000254.

28 S. Zhang, D. Skinner, P. Joshi, E. Criado-Hidalgo, Y.-T. Yeh, J.C. Lasheras, C.R. Caffrey and J.C. del Alamo, J. R. Soc. Interface, 2019, 16, 20180675.

29 S.R. Smithers, R.J. Terry, Parasitology, 1965, 55(4), 695–700.

30 M. Marxer, K. Ingram, and J. Keiser, Parasites & Vectors, 2012, 5, 165.

31 K.A. Heyries, C.L. Hansen, Lab on a Chip, 2011, 23, 3925–3936.

32 C. Poon, J. Mechanical Behavior of Biomedical Materials, 2022, 126, 105024.

33 T.G. Papaioannou, C. Stefanadis, Hellenic J. Cardiol, 2005, 46, 9–15.

34 J.A. Clegg, Experimental Parasitology, 1965, 16(2), 133–147.

35 P.F. Basch, J. Parasitol., 1981, 67(2), 186–190.

36 S.F. Anisuzzaman et al., Front. Immunol., 2021, 12, 635622.

37 L. da Paixão Siqueira, et al., Acta Trop., 2017, 176, 179–187.

38 V Butcher, P.H.H. Schneeberger, J. Keiser, PLoS Negl Trop Dis., 2021, 15(3), e0009313.

39 S.-K. Park, et al., J Biol Chem., 2019, 294(49), 18873–18880.

40 D. Cioli, L. Pica-Mattoccia, Parasitology Research, 2003, 90, S3–S9.

41 S.-K. Park, J.S. Marchant, Trends in Parasitology, 2020, 36(2), 182–194.

42 J.T. Moreira-Filho, A.C. Silva, R.F. Dantas, B.F. Gomes, L.R. Souza Neto, J. Brandao-Neto, R.J. Owens, et al., Front. Immunol., 2021, 12, 642383.

43 M.M. Modena, et al., IEEE Biomedical Circuits and Systems Conference (BioCAS), 2017, 1–4.

44 R. Nacif-Pimenta, A. da Silva Orfano, I.A. Mosley, S.E. Karinshak, et al., Scientific Report, 2019, 9, 10731.

45 S.K. Gokce, E.M. Hegarty, S. Mondal, P. Zhao, N. Ghorashian, et al., Sci. Rep., 2017, 7(1), Art. n^o^ 1.

46 J.C. Weeks, W.M. Roberts, K.J. Robinson, M. Keaney, et al., Int. J. Parasitol. Drugs Drug Resist., 2016, 6(3), 314–328.

47 M.M. Nigo, G.B. Salieb-Beugelaar, M. Battegay, P. Odermatt, P. Hunziker, et al., Prec. Nanomed., 2020, 3(1), 439–458.

48 H.S. Brown, M. Halliwell, M. Qamar, A.E. Read, J.M. Evans, P.N. Wells, Gut, 1989, 30, 503–509.

49 J. Sun, C. Li, S. Wang, Sci Rep., 2016, 6, 34463.

50 J. Sun, W. Hu, C. Li, Parasitol Res., 2013, 112(8), 2983–2990.

51 M.E.M. Saeed, S. Krishna, H.J. Greten, P. G. Kremsner, T. Efferth, et al., Pharmacol Res., 2016, 110, 216–226.

52 E. Bueding, J Gen Physiol., 1950, 33(5), 475–495.

53 A.M. Horemans, A.G. Tielens, S.G. van den Bergh, Mol. Biochem. Parasitol., 1992, 51, 73–79.

54 Z. Faghiri, S.M.R. Camargo, K. Huggel, I.C. Forster, D. Ndegwa, F. Verrey, P.J. Skelly, PLoS One, 2010, 5:e10451.

55 S. Howe, D. Zöphel, H. Subbaraman, C. Unger, J. Held, T. Engleitner, W.H. Hoffmann, A. Kreidenweiss, Antimicrob Agents Chemother., 2015, 59(2), 1193–1199.

56 B.P. Mahoney, N. Raghunand, B. Baggett, R.J. Gillies, Biochem. Pharmacol., 2003, 66(7), 1207–1218.

