## Supplemental figures for "A Self-Purifying Microfluidic System for Identifying Drugs Acting Against Adult Schistosomes"

---

[a.] *Univ. Lille, CNRS, Inserm, CHU Lille, UMR9020-U1277 – CANTHER – Cancer Heterogeneity Plasticity and Resistance to Therapies, Lille F-59000, France.;*

[b.] *CNRS, Univ. Tokyo, IRL2820 - LIMMS, Lille F-59000, France.*

[c.] *Univ. Lille, CNRS, Inserm, CHU Lille, Institut Pasteur de Lille, U1019-UMR9017 - CIIL - Center for Infection and Immunity of Lille, F-59000 Lille, France.*

[d.] *Univ. Lille, CNRS, UPHF, JUNIA, CLI, UMR 8520 – IEMN - Institut d'Electronique, de Microélectronique et de Nanotechnologie, Villeneuve d'Ascq F-59650, France*

† Both authors contributed equally to the work.

* corresponding authors

**Supplementary Figure S1**: **Picture of the single channel chip set-up.** The medium reservoir tubes, the camera, the microfluidic chips, the peristaltic pump are installed onto the hot plate.

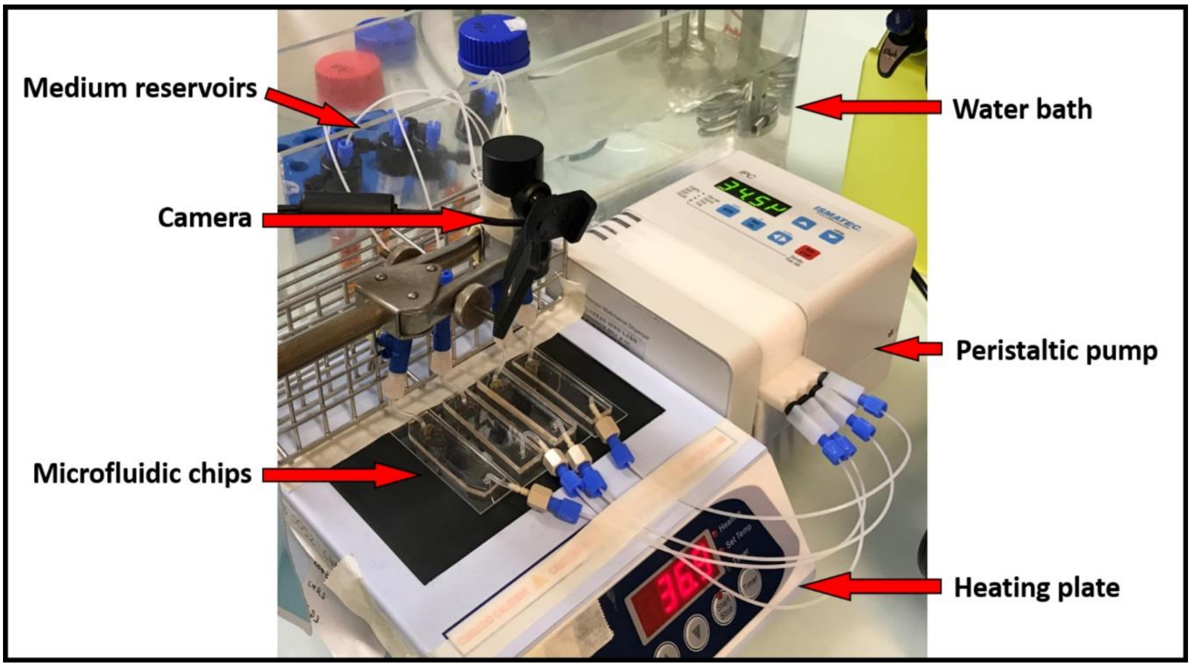

**Supplementary Figure S2**: **Configuration of the microfluidic set-up at the beginning of the experiments for safe introduction of the worms into the device.** Top: view of the entire system with the culture medium reservoirs, the peristaltic pump (ISMATEC, IPC-N8 ISM936 Wertheim, Germany) and the microfluidic chips. Bottom: Zoom on the microfluidic chips that are verticalized to avoid worm clogging in the worm stock and improve worm repartition at the entrance of the microfluidic chip.

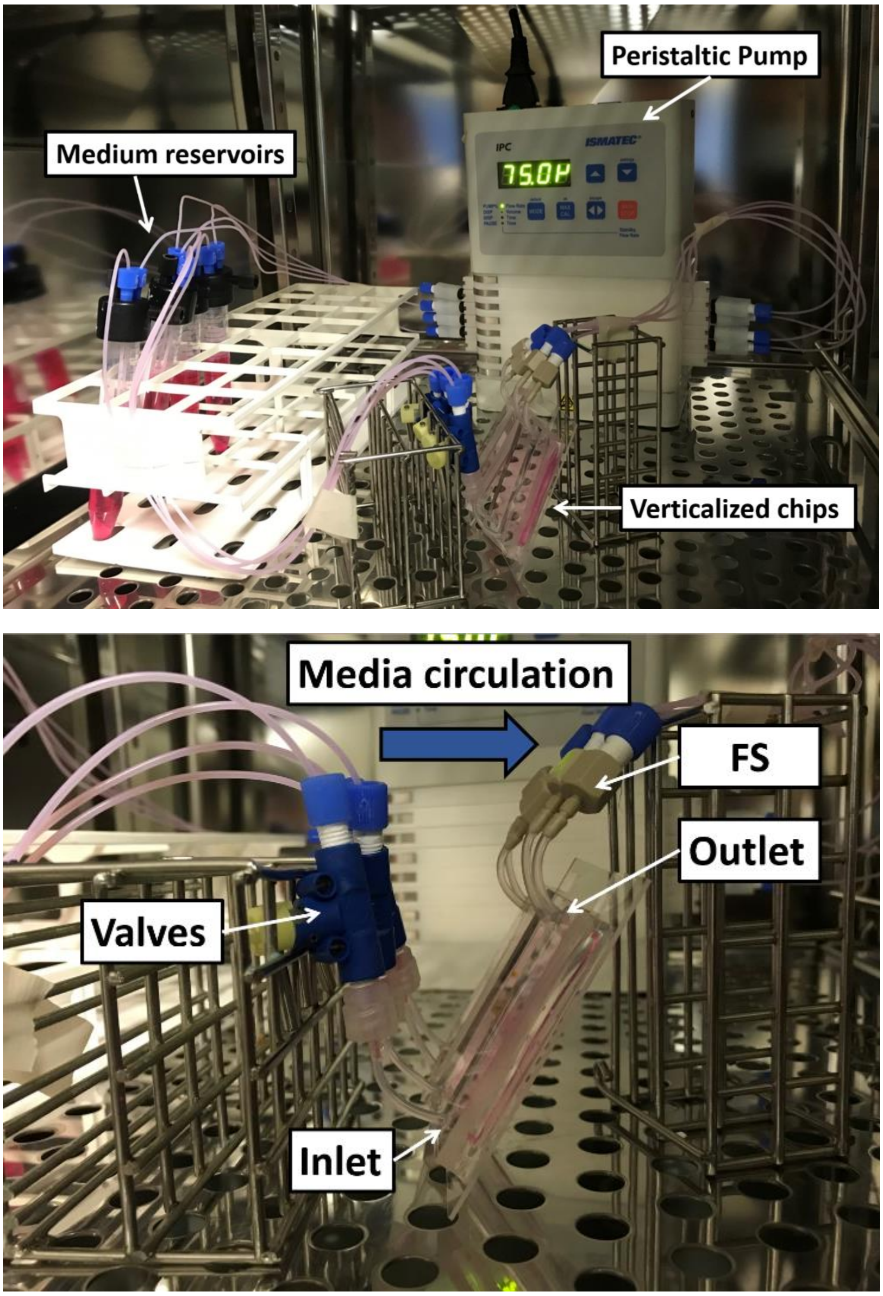

**Supplementary Figure S3**: Progression of the survival rate of Schistosome pairs as a function of time for different culture media (a, RPMI; b, DMEM; c, M199) supplemented or not with serum human, horse or calf in static petri dish culture conditions.

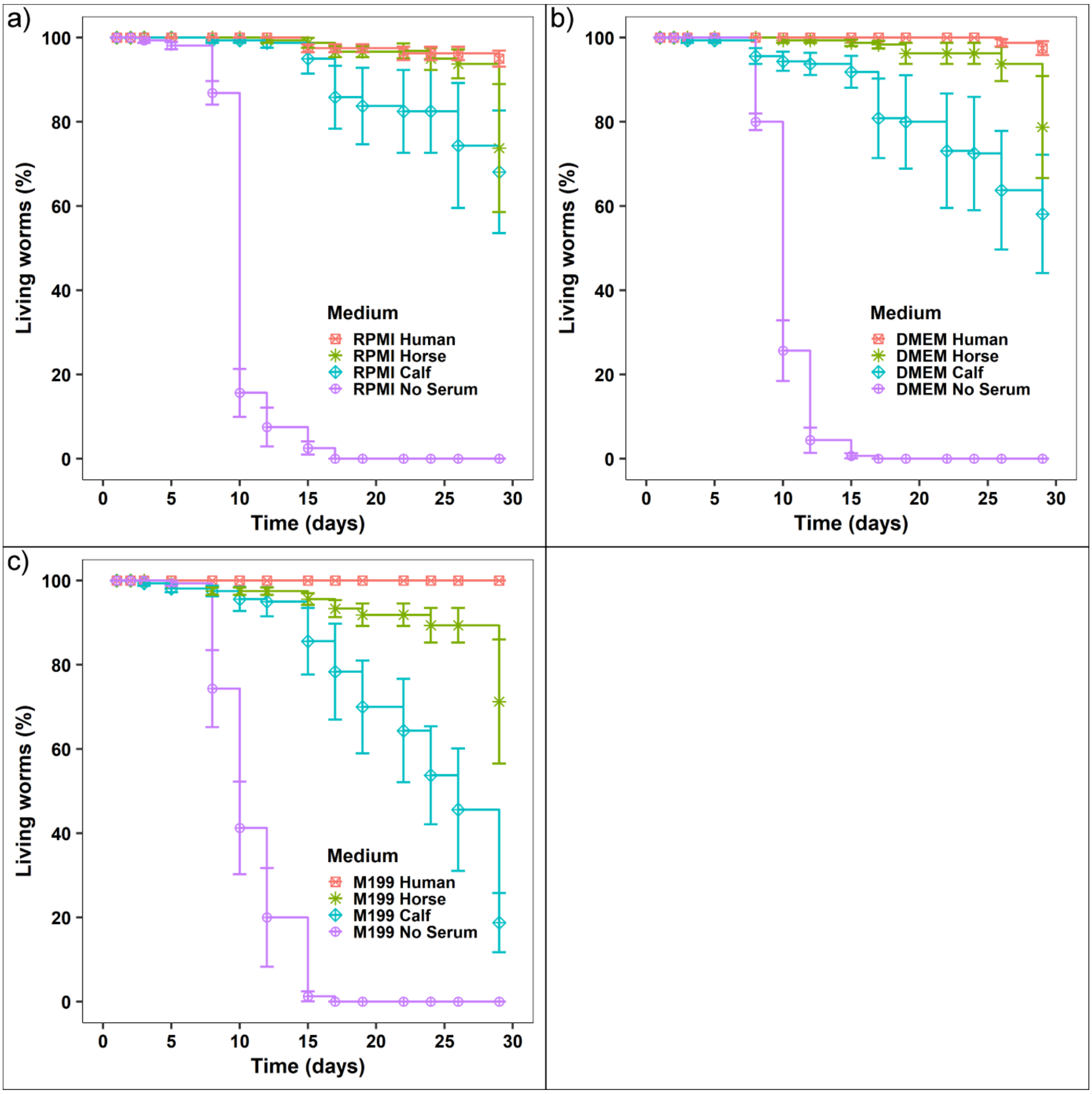

**Supplementary Figure S4**: Percentage of attached worms as a function of flow rates (mL.min$^{-1}$) for the different coatings in the microfluidic device after determination of 2 linear segments based on Table S1 parameters.

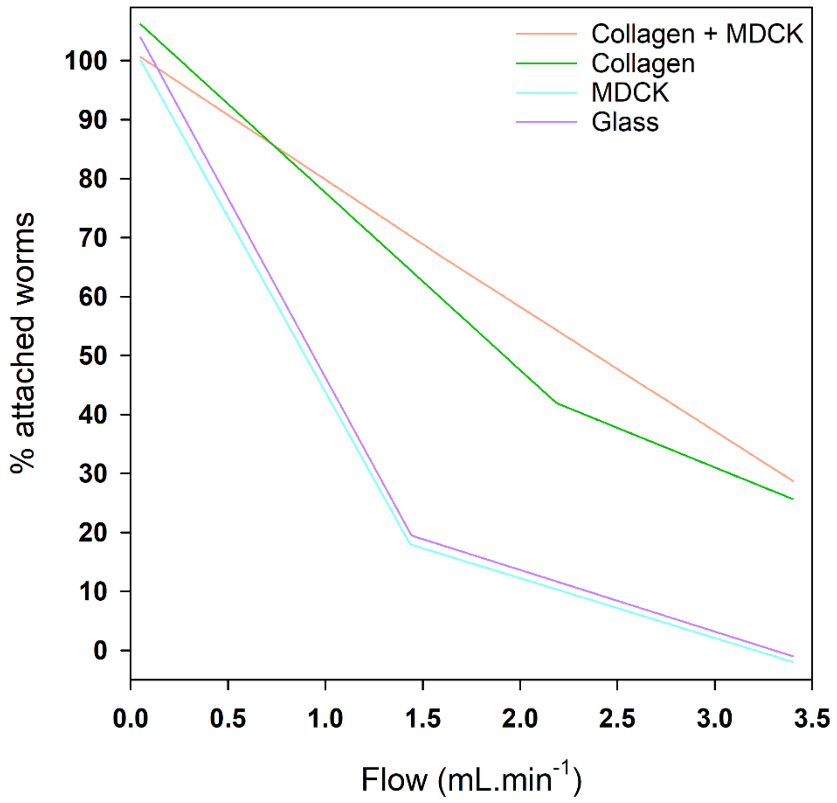

**Supplementary Figure S5**: **Percentage of worm pairs attached to the device as a function of flow rate (mL.min$^{-1}$) for different coatings of the microfluidic device**. Colored lines correspond to the linear regression obtained on the first data segments according to table S2 parameters.

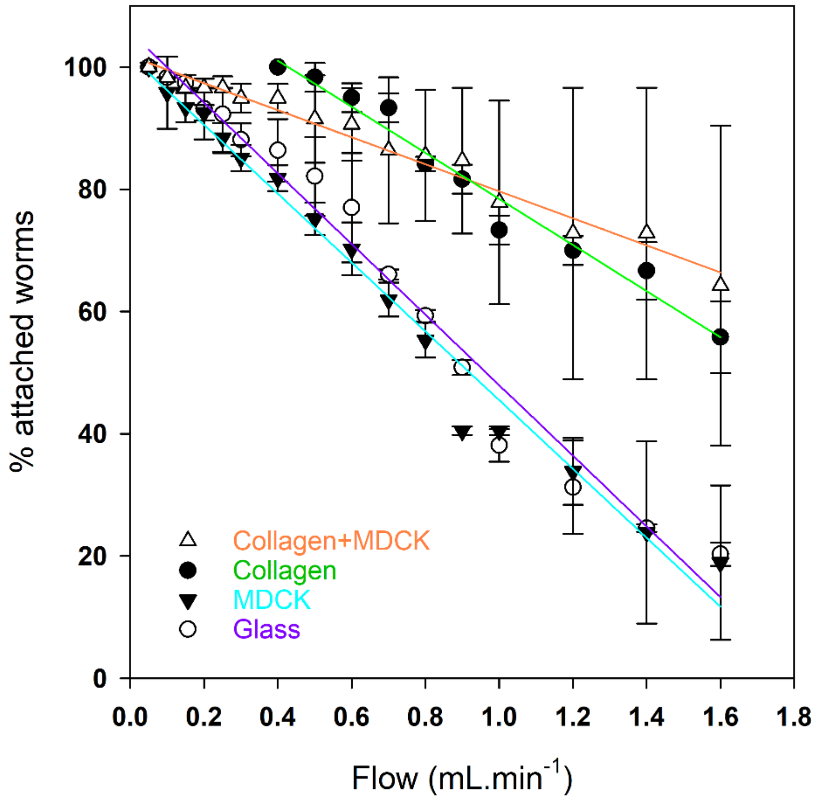

**Supplementary Figure S6:** XY cross-sections of the microfluidic device showing the results of COMSOL simulations for a 1.00 $mL.min^{-1}$ inlet flowrate: a) current line, b) mechanical pressure (Pa), c) culture medium velocity ($m.s^{-1}$).

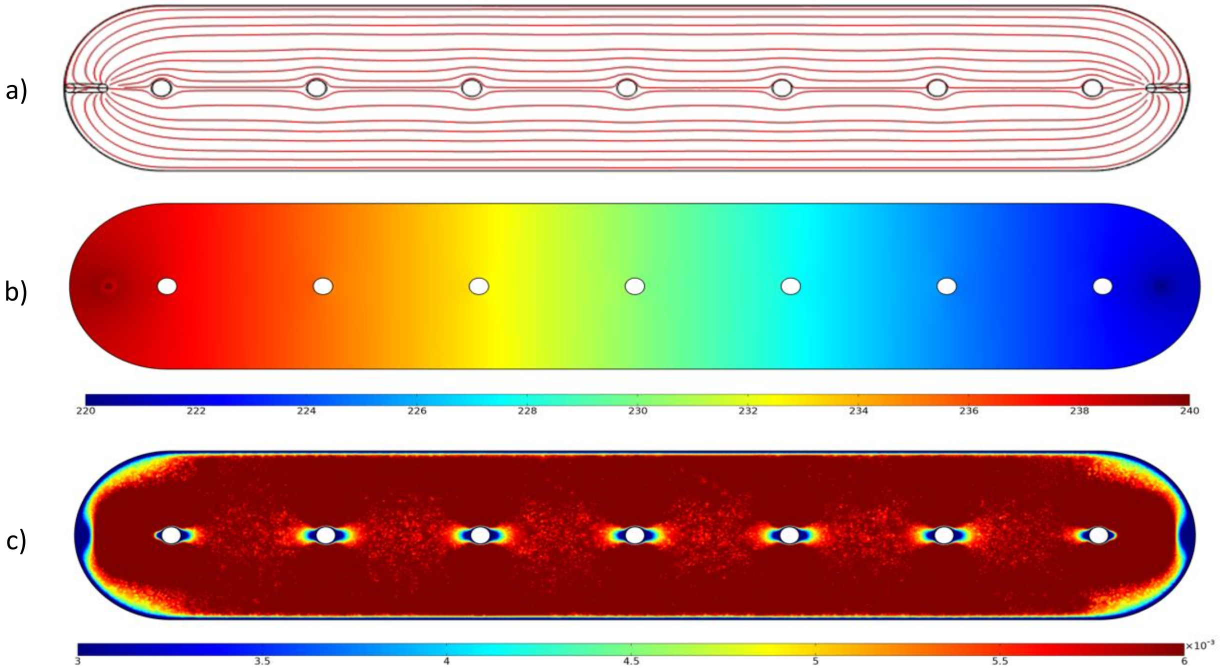

#### Supplementary Table S1

**Calculation of the Transition values (T1) based on the biphasic response observed for worm gripping on glass coated with collagen, MDCK cells or both.** Data have been processed using “Piecewise – linear 2 segments” fitting to obtain first and second segments parameters (Y1, Y2; T1). Next, the corresponding “linear regression” parameters and slope were calculated. Both fittings were performed using SigmaPlot.

| 2 segments | Coll+MDCK | Coll | Glass+MDCK | Glass |
|---|---|---|---|---|
| Y1 | 100.6 | 106.2 | 100.1 | 103.9 |
| SE Y1 | 0.8 | 1.3 | 1.3 | 1.4 |
| Y2 | 66.7 | 41.9 | 18.0 | 19.5 |
| SE Y2 | 46.7 | 4.7 | 2.5 | 2.7 |
| T1 | **1.6** | **2.2** | **1.4** | **1.4** |
| SE T1 | 2.2 | 0.2 | 0.1 | 0.1 |
| slope | -21.2 | -29.4 | -57.2 | -58.6 |

**Supplementary Table S2:**

**Determination of the detachment slope.** Using a linear regression on the first segment data (Y1; T1) on glass coated with collagen, MDCK cells or both, the slopes of worm detachment (in bold) were calculated using SigmaPlot.

| 1st segment | | | | |
|---|---|---|---|---|
| y0 | 101.8 | 116.1 | 101.8 | 105.7 |
| Slope | **-22.1** | **-37.7** | **-56.3** | **-57.8** |
| n sampling | 16 | 10 | 16 | 16 |
| n-2 | 14 | 8 | 14 | 14 |
| SE Slope | 0.9 | 2.3 | 2.1 | 2.4 |
| t/sqr(n-2) | 1.76 | 1.86 | 1.76 | 1.76 |
| 95%Conf | 1.66 | 4.27 | 3.69 | 4.18 |

**Supplementary Table S3:**

**Calculated flow properties in the microfluidic device from COMSOL simulations for different flow rates.**

| **Flow rate (mL.min$^{-1}$)** | **0.05** | **0.30** | **1.00** | **3.40.** |
|---|---|---|---|---|
| Wall shear stress (Pa) | 0,002857 | 0,017179 | 0,058039 | 0,206380 |
| Mean velocity (m.s$^{-1}$) | 0,000312 | 0,001871 | 0,006233 | 0,021129 |
| Reynolds number | 0,13 | 0,79 | 2,62 | 8,88 |
| Pressure gradient (Pa.m$^{-1}$) | 15,9375 | 95,7083 | 319,3750 | 1089,7917 |

**Supplementary Data S1:** Details about the calculation of IC50, IC90 and IC95

Using RStudio, we calculate the percentage of worms in the microfluidic chip as a function of time. 100% corresponds to the number of worms at time where the drug (PZQ or ART) is introduced in the chip, as well as mean and standard deviation values.

We use SIGMAPLOT (SYSTAT Software) to plot the data (percentage of living/attached worms) versus drug concentrations (X) at different time for the microfluidic experiment and at 120h for the static one and used the following fitting equation for the IC50 determination.

$$Y = min - \frac{max - min}{1 + \left(\frac{X}{IC_{50}}\right)^{-hillslope}}$$

| Parameter Values | 5 min | 20 min | 30 min | 50 min |
|---|---|---|---|---|
| **min** | 1.857999099 | 0.38752762 | 1.002628697 | 0.770499143 |
| **max** | 92.83254417 | 82.21514026 | 75.60146965 | 70.00843986 |
| **logEC50** | -6.963866544 | -7.172732304 | -7.214779387 | -7.242491569 |
| **Hillslope** | -3.847098996 | -4.057950592 | -4.654073612 | -5.018832345 |
| **EC50** | 1.09E-07 | 6.72E-08 | 6.10E-08 | 5.72E-08 |
| **Std. Errors** | | | | |
| **min** | 4.422349633 | 3.52285325 | 3.267393583 | 2.738734746 |
| **max** | 3.300897514 | 4.057439538 | 4.007209143 | 3.500095404 |
| **logEC50** | 0.026601346 | 0.03073713 | 0.030699051 | 0.026898542 |
| **Hillslope** | 0.883744748 | 0.864528196 | 1.127693211 | 1.252636418 |
| **95% Confidence Intervals** | | | | |
| **min** | -7,025 to 10,74 | -6,688 to 7,464 | -5,56 to 7,566 | -4,731 to 6,272 |
| **max** | 86,2 to 99,46 | 74,07 to 90,36 | 67,55 to 83,65 | 62,98 to 77,04 |
| **logEC50** | -7,017 to -6,91 | -7,234 to -7,111 | -7,276 to -7,153 | -7,297 to -7,188 |
| **Hillslope** | -5,622 to -2,072 | -5,794 to -2,321 | -6,919 to -2,389 | -7,535 to -2,503 |
| **EC50** | 9,610e-8 to 1,229e-7 | 5,828e-8 to 7,745e-8 | 5,291e-8 to 7,029e-8 | 5,052e-8 to 6,479e-8 |
| **Goodness of Fit** | | | | |
| **Degrees of Freedom** | 50 | 50 | 50 | 50 |
| **R2** | 0.890674603 | 0.867011117 | 0.850097191 | 0.865854099 |
| **Residual Sum of Squares** | 9135.068683 | 9725.977461 | 9592.535516 | 7294.806569 |
| **Sy.x** | 13.51670721 | 13.94702654 | 13.85101839 | 12.07874709 |
| **Fit Status** | Converged | Converged | Converged | Converged |
| **Data** | | | | |
| **Number of X Values** | 10 | 10 | 10 | 10 |
| **Number of Y Replicates** | 11 | 11 | 11 | 11 |
| **Total Number of Y Values** | 54 | 54 | 54 | 54 |
| **Number of Missing Values** | 56 | 56 | 56 | 56 |

ART in fluidic system

| Parameter Values | 5 min | 20 min | 30 min | 50 min |
|---|---|---|---|---|
| min | -8.570641148 | 0.207967993 | -0.091163157 | 0.188956313 |
| max | 95.93999427 | 84.46284733 | 78.42204752 | 72.36317617 |
| logEC50 | -3.857817631 | -4.236874467 | -4.238702437 | -4.237125191 |
| Hillslope | -2.029997902 | -7.125591378 | -7.915683219 | -8.958967093 |
| EC50 | 1.39E-04 | 5.80E-05 | 5.77E-05 | 5.79E-05 |
| **Std. Errors** | | | | |
| min | 16.33724382 | 2.872060057 | 2.614997559 | 2.681727999 |
| max | 3.19625143 | 2.38033321 | 2.195441832 | 2.269340367 |
| logEC50 | 0.07897583 | 0.019091501 | 0.019203185 | 0.022942222 |
| Hillslope | 0.505479475 | 1.533393224 | 1.868533132 | 2.66021107 |
| **95% Confidence Intervals** | | | | |
| min | -41,54 to 24,4 | -5,588 to 6,004 | -5,369 to 5,186 | -5,223 to 5,601 |
| max | 89,49 to 102,4 | 79,66 to 89,27 | 73,99 to 82,85 | 67,78 to 76,94 |
| logEC50 | -4,017 to -3,698 | -4,275 to -4,198 | -4,277 to -4,2 | -4,283 to -4,191 |
| Hillslope | -3,05 to -1,01 | -10,22 to -4,031 | -11,69 to -4,145 | -14,33 to -3,59 |
| EC50 | 9,612e-5 to 0,0002002 | 5,304e-5 to 6,334e-5 | 5,279e-5 to 6,310e-5 | 5,207e-5 to 6,444e-5 |
| **Goodness of Fit** | | | | |
| Degrees of Freedom | 42 | 42 | 42 | 42 |
| R2 | 0.828731666 | 0.93949978 | 0.940561615 | 0.925711211 |
| Residual Sum of Squares | 6280.112266 | 4433.037685 | 3858.822155 | 4236.275845 |
| Sy.x | 12.22810216 | 10.27368076 | 9.585234373 | 10.04309268 |
| Fit Status | Converged | Converged | Converged | Converged |
| **Data** | | | | |
| Number of X Values | 10 | 10 | 10 | 10 |
| Number of Y Replicates | 9 | 9 | 9 | 9 |
| Total Number of Y Values | 46 | 46 | 46 | 46 |
| Number of Missing Values | 44 | 44 | 44 | 44 |

### PZQ in static system

| Parameter Values | Horse serum | Calf serum |
|---|---|---|
| min | -1.70E-09 | -5.41E-10 |
| max | 100 | 100 |
| logEC50 | -6.610438254 | -6.610838208 |
| Hillslope | -49.29984351 | -119.197851 |
| EC50 | 2.45E-07 | 2.45E-07 |
| Std. Errors | | |
| min | 1.02E-09 | 3.03E-10 |
| max | 7.25E-10 | 2.56E-10 |
| logEC50 | 3.76E-03 | 8.38E-04 |
| Hillslope | 0.034869311 | 1.59499281 |
| 95% Confidence Intervals | | |
| min | -3,953e-9 to 5,583e-10 | -1,155e-9 to 7,314e-11 |
| max | 100, to 100, | 100, to 100, |
| logEC50 | -6,619 to -6,602 | -6,613 to -6,609 |
| Hillslope | -49,38 to -49,22 | -122,4 to -116, |
| EC50 | 2,406e-7 to 2,499e-7 | 2,440e-7 to 2,460e-7 |
| Goodness of Fit | | |
| Degrees of Freedom | 11 | 38 |
| R2 | 1 | 1 |
| Residual Sum of Squares | 3.47E-17 | 5.25E-17 |
| Sy.x | 1.77E-09 | 1.18E-09 |
| Fit Status | Converged | Did not converge |
| Data | | |
| Number of X Values | 5 | 14 |
| Number of Y Replicates | 3 | 3 |
| Total Number of Y Values | 15 | 42 |
| Number of Missing Values | 0 | 0 |

ART in static conditions

| Parameter Values | Horse serum | Calf serum |
|---|---|---|
| min | -0.121355269 | -9.627717157 |
| max | 100.1067963 | 100.1606246 |
| logEC50 | -3.755743058 | -3.681147199 |
| Hillslope | -4.890928411 | -2.131994726 |
| EC50 | 1.76E-04 | 2.08E-04 |
| Std. Errors | | |
| min | 2.263383942 | 4.313503677 |
| max | 2.023949399 | 2.046973458 |
| logEC50 | 0.034275193 | 0.025716213 |
| Hillslope | 2.238503178 | 0.211067045 |
| 95% Confidence Intervals | | |
| min | -4,976 to 4,733 | -18,36 to -0,8953 |
| max | 95,77 to 104,4 | 96,02 to 104,3 |
| logEC50 | -3,829 to -3,682 | -3,733 to -3,629 |
| Hillslope | -9,692 to -0,08972 | -2,559 to -1,705 |
| EC50 | 0,0001482 to 0,0002079 | 0,0001848 to 0,0002349 |
| Goodness of Fit | | |
| Degrees of Freedom | 14 | 38 |
| R2 | 0.99120117 | 0.977129253 |
| Residual Sum of Squares | 333.4756528 | 1353.744024 |
| Sy.x | 4.880541925 | 5.96865502 |
| Fit Status | Converged | Converged |
| Data | | |
| Number of X Values | 6 | 14 |
| Number of Y Replicates | 3 | 3 |
| Total Number of Y Values | 18 | 42 |
| Number of Missing Values | 0 | 0 |
